# Antisense targeting of decoy exons can reduce intron retention and increase protein expression in human erythroblasts

**DOI:** 10.1101/2020.01.13.902544

**Authors:** Marilyn Parra, Weiguo Zhang, Jonathan Vu, Mark DeWitt, John G. Conboy

## Abstract

The decoy exon model has been proposed to regulate a subset of intron retention (IR) events involving predominantly larger introns (>1kb). Splicing reporter studies have shown that decoy splice sites are essential for activity, suggesting that decoys act by engaging intron-terminal splice sites and competing with cross-intron interactions required for intron excision. The decoy model predicts that antisense oligonucleotides blocking decoy splice sites in endogenous pre-mRNA should increase productive gene expression by reducing IR. Indeed, we now demonstrate that targeting a decoy 5′ splice site in the O-GlcNAc transferase (OGT) gene reduced IR from ∼80% to ∼20% in primary human erythroblasts, accompanied by increases in spliced OGT RNA and OGT protein expression. The remaining OGT IR was refractory to antisense treatment and might be mediated by independent mechanism(s). In contrast, other retained introns were strongly dependent on decoy function, since IR was nearly eliminated by antisense targeting of 5′ splice sites. Genes in the latter group encode the widely expressed splicing factor (SF3B1), and the erythroid-specific structural protein, alpha-spectrin (SPTA1). These results show that modulating decoy exon function can dramatically alter IR, and suggest that dynamic regulation of decoy exons could be a mechanism to fine tune gene expression post-transcriptionally in many cell types.

## Introduction

Gene expression is determined not only by transcription rate, but also by post-transcriptional processes including the efficiency with which pre-mRNA is spliced into translatable mRNA. Intron retention (IR) is a form of RNA processing that selectively modulates splicing of specific introns (Boutz et al. 2015; Braunschweig et al. 2014; Mauger et al. 2016; Jacob and Smith 2017), in essence rendering them ‘alternative introns’. By regulating the efficiency of intron splicing, cells can alter the balance between two competing pathways: one that generates fully spliced mRNA that can be translated into protein, and a second that produces incompletely spliced “intron retention” transcripts (IR-transcripts). Most of the latter contain premature translation termination signals that preclude synthesis of full length protein. IR-transcripts that are otherwise spliced and polyadenylated can experience several fates in different cellular contexts. Such transcripts are often detained in the nucleus, where they may be degraded (Pendleton et al. 2018) or they may serve as a reservoir for new mRNA production via excision of the retained intron(s) (Boothby et al. 2013; Mauger et al. 2016; Ninomiya et al. 2011); in many cases, the fate is unknown. Alternatively, IR transcripts can be exported to the cytoplasm for degradation by nonsense-mediated decay (NMD) (Wong et al. 2013), or they may persist for translation (Rekosh and Hammarskjold 2018) or other unknown functions (Brugiolo et al. 2017). At a constant transcription rate, greater diversion of pre-mRNA into untranslated IR-transcripts should reduce output of mRNA and decrease protein synthesis. Coordinate regulation of IR can effect programmed changes in gene expression patterns during normal development as cells differentiate and respond to environmental signals (Boutz et al. 2015; Ni et al. 2016; Wong et al. 2013; Mauger et al. 2016; Braun et al. 2017; Naro et al. 2017; Edwards et al. 2016; Pimentel et al. 2016; Shalgi et al. 2014). Conversely, aberrations in the IR program are observed in many diseases including cancers where they can adversely impact expression of many genes (Dvinge and Bradley 2015; Luisier et al. 2018; Adusumalli et al. 2019). Although mechanisms of IR are not well understood, RNA binding proteins (RBPs) (Cho et al. 2014; Pendleton et al. 2017) and factors that modify RBPs (Braun et al. 2017) have been shown impact IR. In a few cases RNA sequence elements required for regulating individual intron retention events have been identified (Park et al. 2017; Pendleton et al. 2017; Rekosh and Hammarskjold 2018; Parra M. et al. 2018).

Analysis of RNA-seq profiles from differentiating erythroid cell populations revealed highly dynamic, global changes in the erythroid transcriptome, including changes in RNA processing of both cassette exons and retained introns, as the cells undergo extensive remodeling during the final cell divisions prior to enucleation (An et al. 2014; Pimentel et al. 2014; Pimentel et al. 2016; Edwards et al. 2016). The IR program encompasses hundreds of IR-transcripts that are polyadenylated and spliced except for selective retention of one or more introns (Pimentel et al. 2016; Edwards et al. 2016). In late erythroblasts, numerous IR transcripts are abundantly-expressed, many of which comprise ≥25% of the steady state RNA from their cognate genes. Some of these are dynamically regulated during terminal erythropoiesis, while others exhibit stable IR levels, indicating multiple regulatory pathways (Pimentel et al. 2016). While the majority of erythroblast retained introns are short (<1kb), as observed in other systems (Braunschweig et al. 2014), a subset of important erythroid genes exhibit larger retained introns having embedded decoy exon(s) that are essential for retention (Parra M. et al. 2018). According to the decoy model, cryptic decoy exon(s) interact nonproductively with intron-terminal splice sites, engaging them in a manner that fails to stimulate efficient splicing catalysis. By competing with cross intron interactions necessary for intron removal, decoy interactions promote IR. Supporting evidence for this model includes the ability of decoy exons to activate IR in heterologous splicing reporters; the dependence of this IR activity on intact decoy splice sites; and the enrichment of U2AF binding at 3’ splice sites of decoy exons (Parra M. et al. 2018).

Here we explored the hypothesis that decoy exon function can be modulated in primary erythroid progenitors to alter endogenous RNA processing fates and thereby tune gene expression. Given the ability of antisense oligonucleotides to alter splicing outcomes by masking regulatory elements in deep intron space (Parra M. K. et al. 2012; Lovci et al. 2013; Sibley et al. 2015), we employed a similar strategy to test whether targeting decoy splice sites with antisense reagents can inhibit IR. New results indicate that blocking highly conserved decoy exons in three broadly expressed genes (SF3B1, OGT, and SNRNP70), and in an erythroid-specific gene (SPTA1), greatly reduces intron retention activity in endogenous transcripts, and can increase spliced RNA and protein expression. These results validate the function of decoy exons in the context of their natural endogenous transcripts, and suggest that many of the ∼400 predicted decoys in differentiating human erythroblasts could be regulated to impact protein expression.

## Results

### Decoy exon targeting strategy

Candidate decoy exons were identified in retained introns of NMD-inhibited erythroblasts by virtue of the novel splice junctions created when they splice, albeit inefficiently, to adjacent exons (Parra M. et al. 2018). The decoy model hypothesizes that their main function is to form early spliceosomal complexes with intron-terminal splice sites that become arrested at a pre-catalytic stage of assembly; catalytic splicing at decoy splice sites is inefficient and typically leads to NMD. To assess decoy function in endogenous erythroid transcripts, we reasoned that antisense oligonucleotides targeting decoy exons should interfere with IR to reduce retention efficiency. To maximize our ability to detect such changes, we selected IR-transcripts meeting the following criteria: (1) the transcript must possess a unique intron exhibiting ≥20% retention in late erythroblasts; (2) its cognate gene must be expressed in moderate to high abundance; and (3) the embedded decoy exons must have simple splice site architecture. The last feature served to restrict analysis to decoys that either have unique splice junctions, or have closely spaced alternative junctions that can be blocked with a single 25nt antisense morpholino (MO). This design was expected to maximize the likelihood of blocking spliceosome assembly at the decoy. However, it eliminated from consideration strong decoys in ARGLU1 and DDX39B that possess alternative splice sites distributed over a wider range (Parra M. et al. 2018; Pirnie et al. 2017). Finally, we targeted 5’ splice sites, because the relatively low GC content at typical 3’ splice sites was predicted to reduce MO affinity and effectiveness.

Figure 1A shows relevant features of the IR regions from four genes chosen for analysis. OGT intron 4 (3.3kb), SF3B1 intron 4 (1.8kb), SNRNP70 intron 7 (3.2kb), and SPTA1 intron 20 (1.8kb) all exhibit substantial retention in erythroid progenitors at day 9 of the culture (D9) and in well-differentiated erythroblasts at day 16 (D16). Each of these introns encodes decoy exon(s), not represented in Refseq annotations, that were defined by analysis of splice junction reads (Parra M. et al. 2018) and are depicted in a custom reannotation track (Figure 1A). The decoys in OGT, SF3B1, and SNRNP70 have been highly conserved from fish to mammals, while the SPTA1 decoy is conserved only among mammals. In previous assays with splicing reporters, the OGT and SF3B1 decoys exhibited strong IR activity, while the activity of the SNRNP70 decoy had weaker activity (Parra M. et al. 2018). The SPTA1 decoy has not been assayed previously for IR activity.

**Figure 1.**
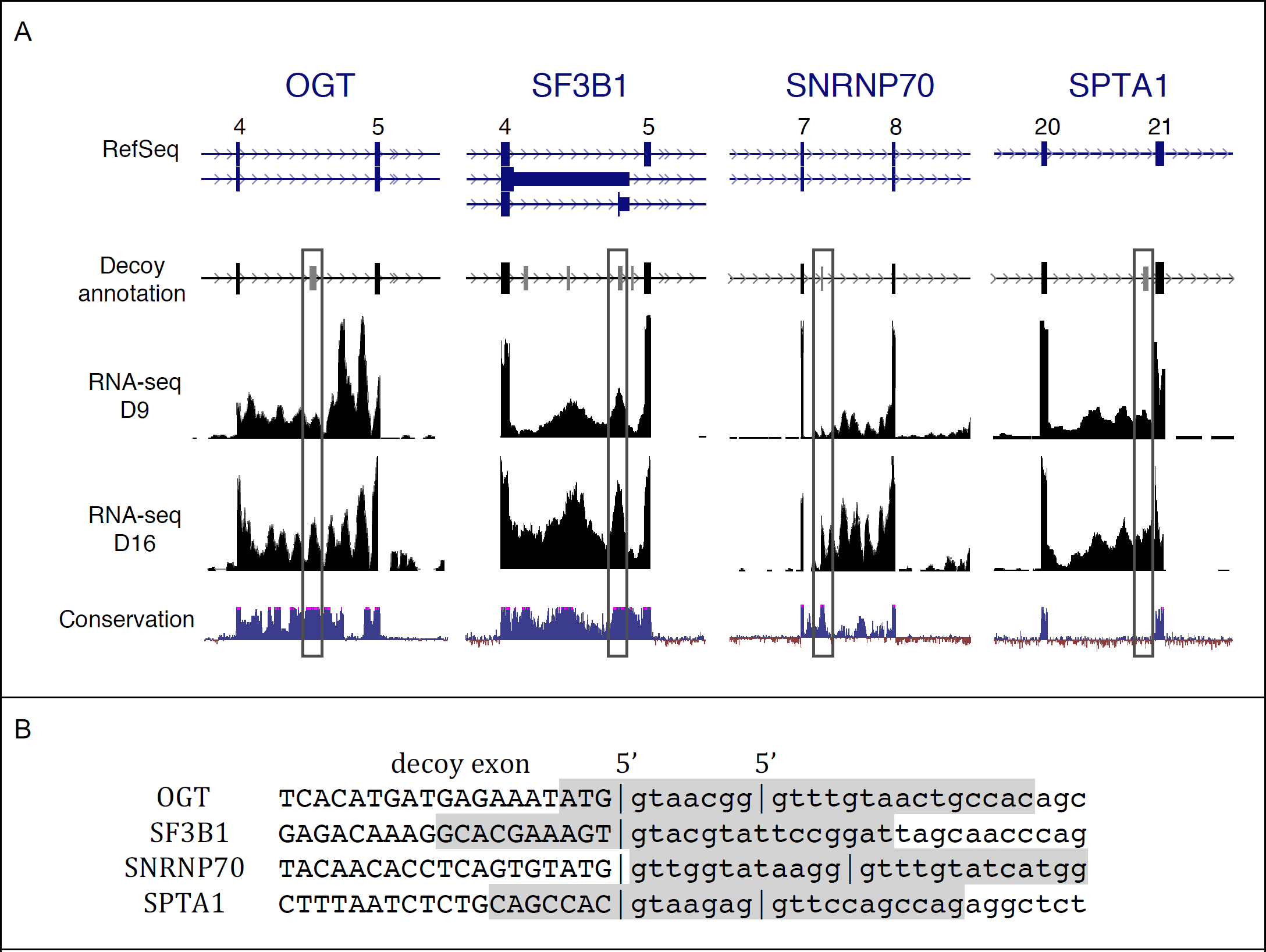
Intron retention and candidate decoy exons in targeted erythroid genes. 1A. Annotation of key features in IR regions of four prominent erythroid genes. Top panel: Refseq gene annotations, lacking indications of intron retention isoforms or decoy exons predicted within the introns. For SF3B1, only the four frequently spliced decoys are shown, from a total six total (Parra M. et al. 2018). Lower panels show a reannotation that includes predicted decoy exons (boxed), RNA-seq data from early stage (D9) and late stage (D15) erythroblasts, and phylogenetic conservation of the relevant gene regions. 1B. 5’ splice site features of targeted decoy exons. Upper case, decoy exon sequence; lower case, downstream intron sequence. Vertical bars show 5’ splice site junctions identified in RNA-seq data from erythroblasts inhibited for nonsense-mediated decay. Shaded regions indicate regions targeted by antisense morpholinos.

The 5’ splice site regions of decoys targeted in this study are shown in Figure 1B. The SF3B1 decoy exhibits only one 5’ splice site, while the other three decoys all have alternative 5’ splice sites located within 7-12nt of each other. The presence of multiple splice sites could be integral to the decoy mechanism, since this appeared to be a frequent feature of decoy exons, and because it has been shown that concurrent occupancy of alternative splice sites can inhibit splicing (Chen et al. 2017). The shaded regions indicate sequences targeted by antisense morpholinos in the IR assays below.

### Reduction of intron retention by antisense targeting of decoy 5’ splice sites

Primary human erythroid cultures were electroporated with antisense morpholinos, then cells were cultured for two days under standard conditions. RNA was then isolated for analysis by RT-PCR to investigate changes in the balance between IR-transcripts and spliced transcripts. We studied the effects of decoy targeting in four different genes using this approach. The targeting scheme and PCR strategy for analysis of IR in the OGT gene, which encodes O-GlcNAC transferase, is shown in Figures 2A. The decoy in OGT intron 4 exhibited strong IR activity in minigene splicing reporters (Parra M. et al. 2018). In endogenous OGT transcripts, we first assessed retention of the full length intron 4 by standard RT-PCR analysis under conditions that interrogate the E3-E6 region. Control cells treated with an irrelevant MO yielded two major OGT amplification products (Figure 2B, lane 1): a short product representing spliced mRNA, and a larger product corresponding to an IR transcript in which introns 3 and 5 were removed but intron 4 retained. Cells treated with the OGT decoy-specific MO exhibited a substantial decrease in the IR isoform (Figure 2B, lane 2). In contrast, the OGT MO did not alter retention of a heterologous decoy-containing intron in the SF3B1 gene, confirming specificity of the MO effects on IR (Figure 2C, compare lanes 1 and 2). These results strongly support the hypothesis that full length introns are specifically retained in a subset of transcripts, and that retention can be greatly suppressed by anti-decoy MOs.

**Figure 2.**
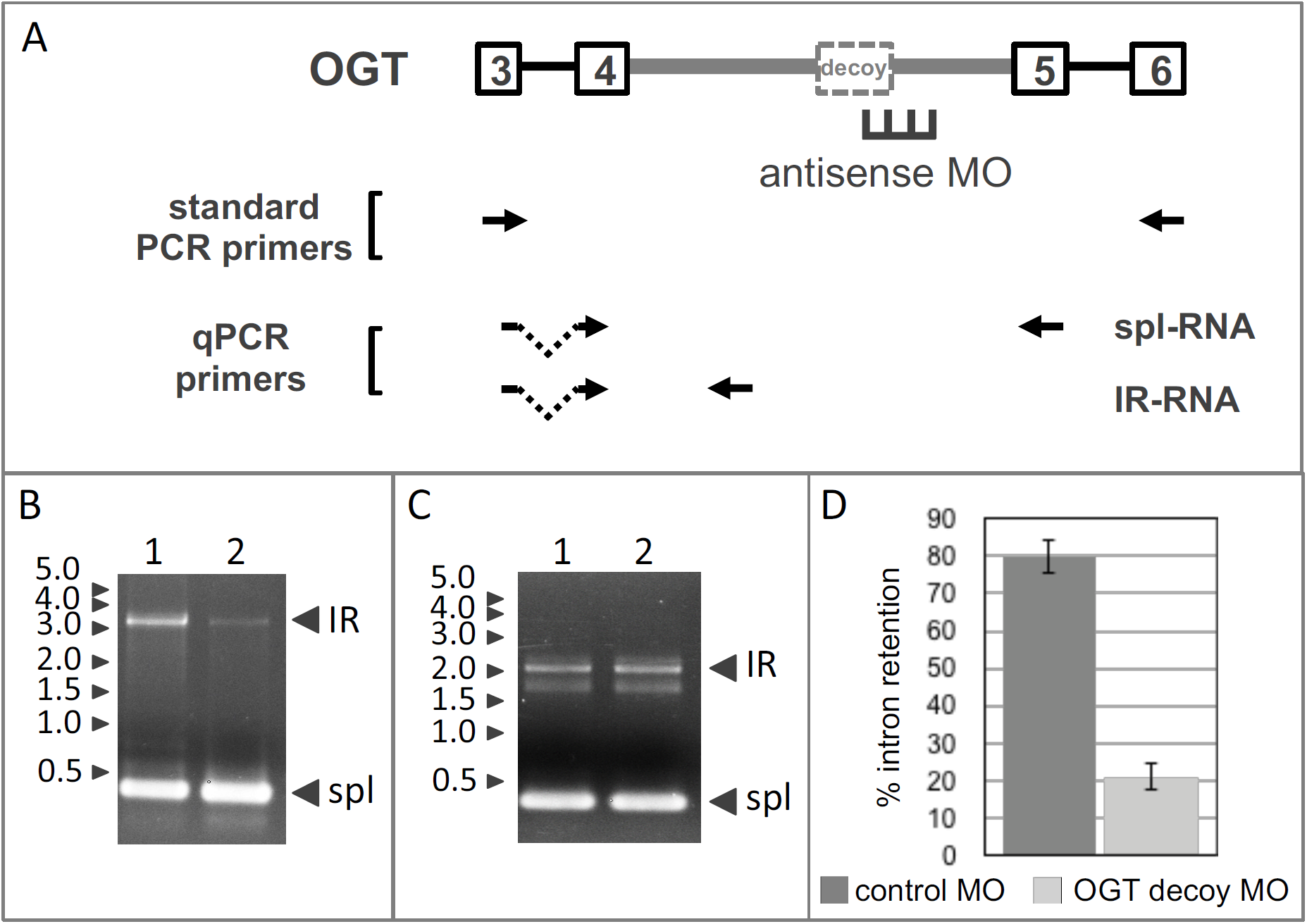
Antisense inhibition of OGR IR. A. OGT gene structure in the IR region showing retained intron 4 (thick gray line) with its decoy exon and flanking exons. Position of antisense MO designed to block the 5’ splice site is shown, along with primer pairs used for RT-PCR. B. Gel analysis of IR and spliced bands amplified from endogenous OGT transcripts by standard RT-PCR from cells treated with negative control MO (lane 1) or OGT decoy-specific MO (lane 2). C. As a control for decoy-specific effects of the MO treatment, gel analysis of IR and spliced bands amplified from endogenous SF3B1 transcripts after treatment with negative control MO (lane 1) or OGT decoy-specific MO (lane 2). D. OGT IR, as a percentage of total OGT transcripts, in cells treated with negative control or decoy-specific MO. IR was assessed using RT-qPCR. Results show average IR from data of four experiments. Error bars indicate standard deviation.

However, standard RT-PCR does not provide a quantitative measure of PIR (percent intron retention), in part due to inefficient amplification of long retained introns. We therefore performed RT-qPCR using primers that amplify unique regions of the IR isoforms or the spliced isoforms, respectively. For OGT, the fraction of transcripts bearing the retained intron was estimated at ∼80% in control cells, but PIR was substantially reduced to ∼21% in cells targeted with the OGT decoy 5’ splice site MO. Interestingly, the level of OGT IR did not decrease further when the MO concentration was doubled (results not shown), suggesting that a component of OGT IR is modulated in a decoy-independent manner.

The next decoy-mediated IR event selected for analysis was in the SF3B1 gene (Figure 3A). We focused on decoy exon 4e, shown previously to exhibit the strongest IR activity among several potential decoys in SF3B1 intron 4 (Parra M. et al. 2018). Similar to OGT, cells treated with the SF3B1-specific MO exhibited much-reduced amounts of the IR-transcript when examined by standard RT-PCR (Figure 3B, compare lanes 1 and 2). Quantitation by qPCR yielded a different result than was observed for OGT, since PIR in controls cells (∼26%) was almost eliminated by the SF3B1 decoy 5’ splice site MO (∼3%).

**Figure 3.**
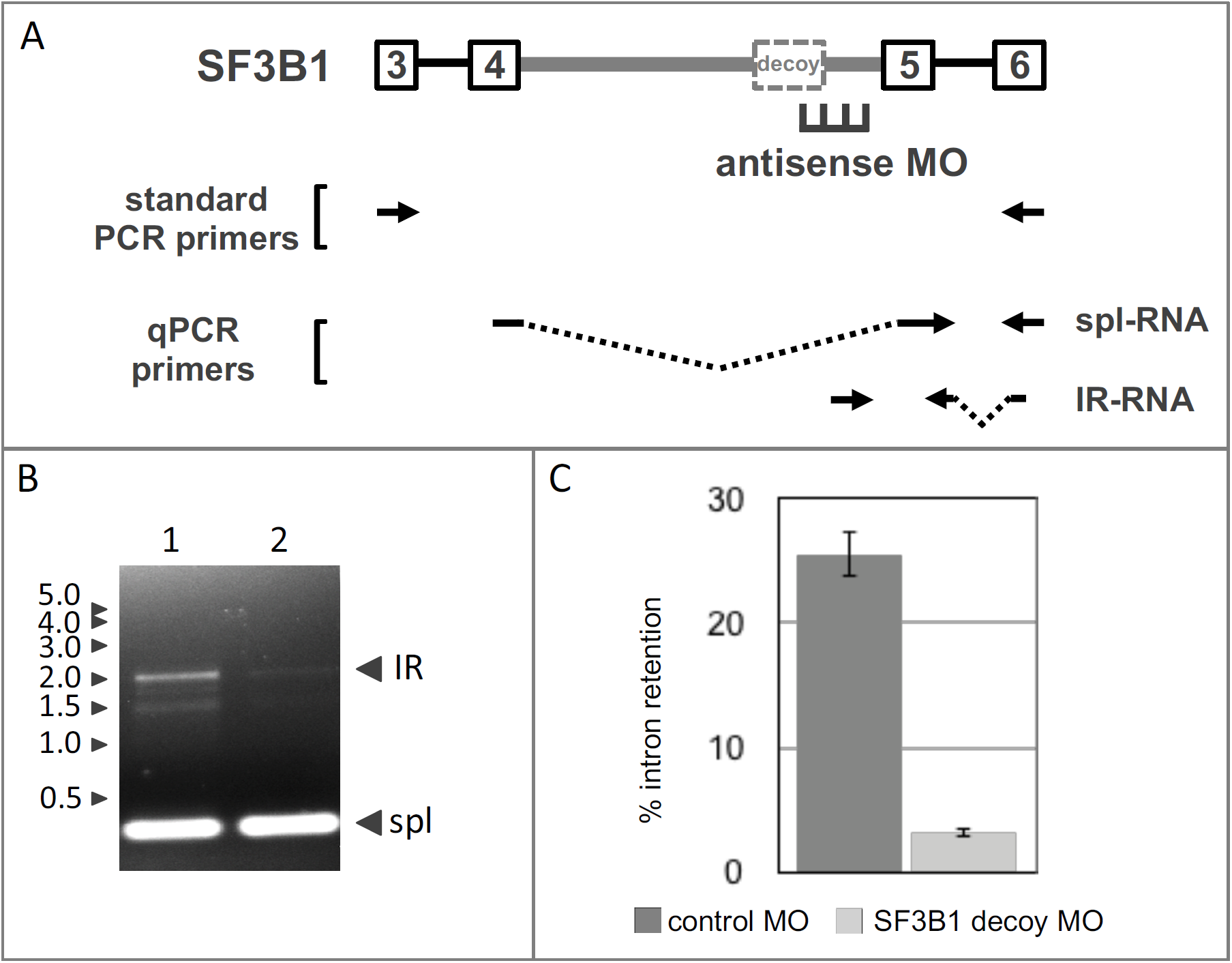
Antisense inhibition of SF3B1 IR. A. SF3B1 gene structure in the IR region showing retained intron 4 (thick gray line) and its major decoy exon, together with adjacent introns and exons. Position of antisense MO designed to block the 5’ splice site is shown, along with primer pairs used for RT-PCR. B. Gel analysis of IR and spliced bands amplified from endogenous SF3B1 transcripts by standard RT-PCR from cells treated with negative control MO (lane 1) or SF3B1 decoy-specific MO (lane 2). C. SF3B1 IR, as a percentage of total SF3B1 transcripts, in cells treated with negative control or decoy-specific MO. IR was assessed using RT-qPCR to compare the relative amounts of IR and spliced products. Results show average IR of three experiments. Error bars indicate standard deviation.

The two remaining targets represented decoys about which less prior information was known than for OGT and SF3B1. The predicted decoy exon in SNRNP70 is 60/72nt, depending on alternative 5’ splice site choice, and might have unique properties since retention has been observed primarily only for downstream intron sequences (Figure 1A). Moreover, this decoy exhibited only weak IR activity in a heterologous splicing reporter (Parra M. et al. 2018). For SPTA1, an 80/87nt noncoding decoy exon mapping near the 3’ end of retained intron 20 was predicted on the basis of splice junction reads. A few RNA-seq reads spanned the SPTA1 decoy exon and linked it to both exon 19 upstream and exon 20 downstream, confirming its potential to be spliced at low frequency (data not shown). Given that SPTA1 encodes an abundant and erythroid-specific structural protein, alpha spectrin, control of IR could be important in regulating assembly of the erythroid membrane skeleton during terminal erythropoiesis.

Figures 4A and 4B show the targeting approach and PCR strategies used to test IR-promoting activity for predicted decoy exons in SNRNP70 and SPTA1. The effects of decoy-specific antisense MOs were assessed by RT-qPCR to quantitate both IR transcripts and fully spliced transcripts (Figures 4C and 4D). Electroporation of human erythroblasts with a MO against the 5’ splice site region of SNRNP70’s decoy substantially reduced the level of IR from about 35% in control erythroid cells to about 9% in MO-treated cells (Figure 4C). Interestingly, the SPTA1 decoy-specific MO also strongly inhibited IR, from ∼20% down to only ∼2%. Together these results strongly support the hypothesis that decoy exons represent a novel regulatory component of the gene expression program, in which they can quantitatively modulate mRNA expression levels by tuning the splicing efficiency of key retained introns.

**Figure 4.**
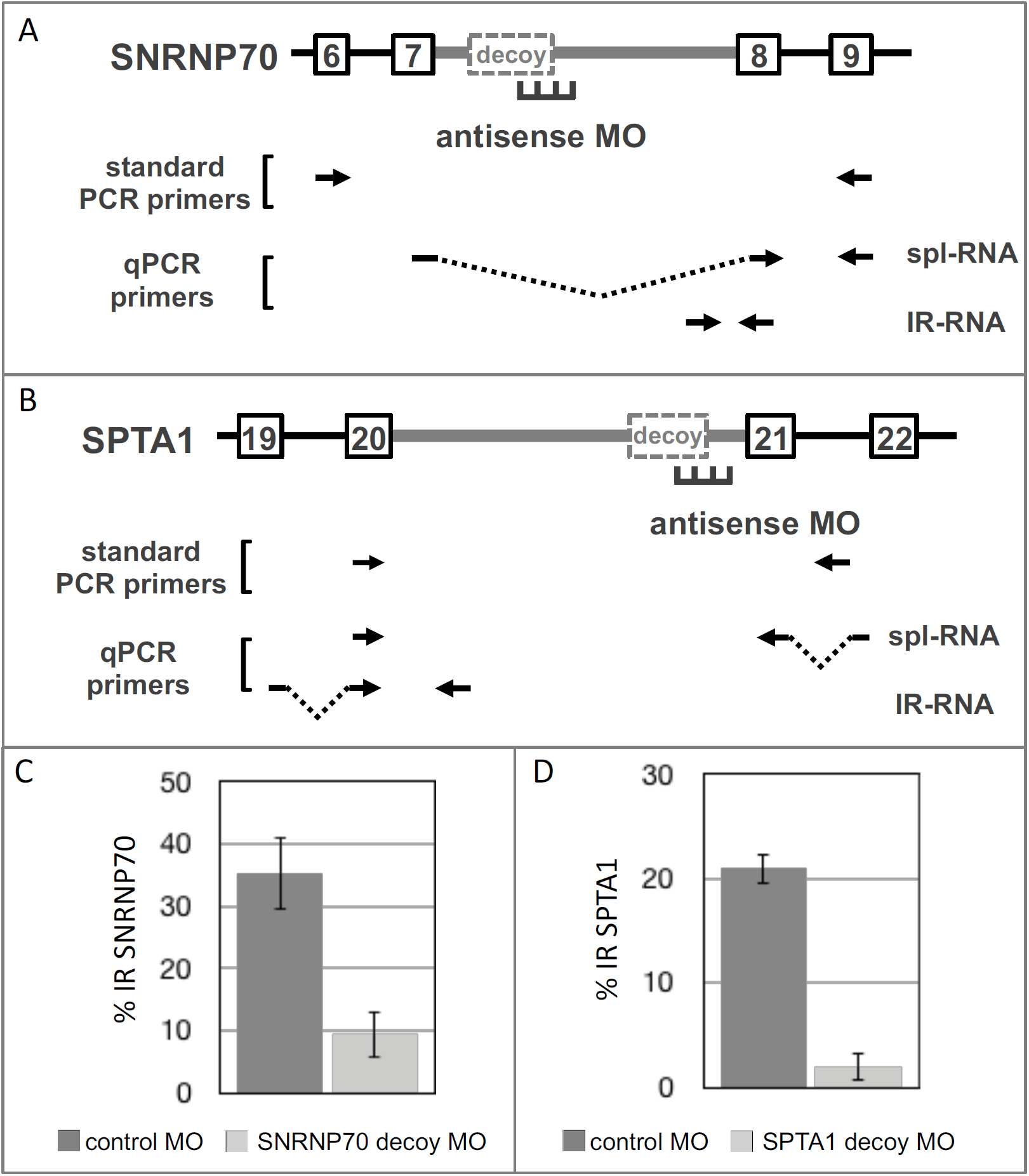
Antisense inhibition of SNRNP70 IR and SPTA1 IR. A. SNRNP70 gene structure in the IR region showing retained intron 7 (thick gray line) with its major decoy exon, together with adjacent introns and exons. Position of antisense MO designed to block the 5’ splice site is shown, along with primer pairs used for RT-PCR. B. SPTA1 gene structure in the IR region showing retained intron 20 (thick gray line) with its major decoy exon, together with adjacent introns and exons. Position of antisense MO designed to block the 5’ splice site is shown, along with primer pairs used for RT-PCR. C and D. IR in cells treated with SNRNP70-specific (C) or SPTA1-specific (D) MO, in parallel with cells subjected to control MO treatment. IR was assessed using RT-qPCR to compare the relative amounts of IR and spliced products.

### Impact of decoy targeting on expression of spliced RNA and protein

The dramatic reduction in IR for several genes suggests that inhibition of decoy exon function should lead to increased expression of spliced mRNA and increased capacity for protein synthesis. We explored this issue using OGT as a model, since the large MO-induced reduction in PIR would be expected to yield a significant increase in protein expression. Based on the 4-fold difference in IR between control cells and cells treated with the OGT decoy-specific MO, measured at 48hrs post-electroporation, one might predict a similar 4-fold increase in OGT mRNA and protein. Analysis of qPCR data revealed that the spliced OGT transcripts were actually increased 1.7-2.7-fold, when normalized to actin transcript expression in the same cells (Figure 5A). To explore the reason for the modest discrepancy in expected vs observed expression, we quantitated total OGT transcript levels (spliced plus IR transcripts) and found that overall OGT RNA expression was reduced in comparison to control cells. Therefore, the 4-fold increase in splicing efficiency was partially offset by reduced steady state OGT RNA levels, presumably due to other compensatory mechanisms as part of O-GlcNAc homeostasis. Nevertheless, this result confirms that regulation of decoy-mediated IR effected a significant change in spliced OGT RNA expression.

**Figure 5.**
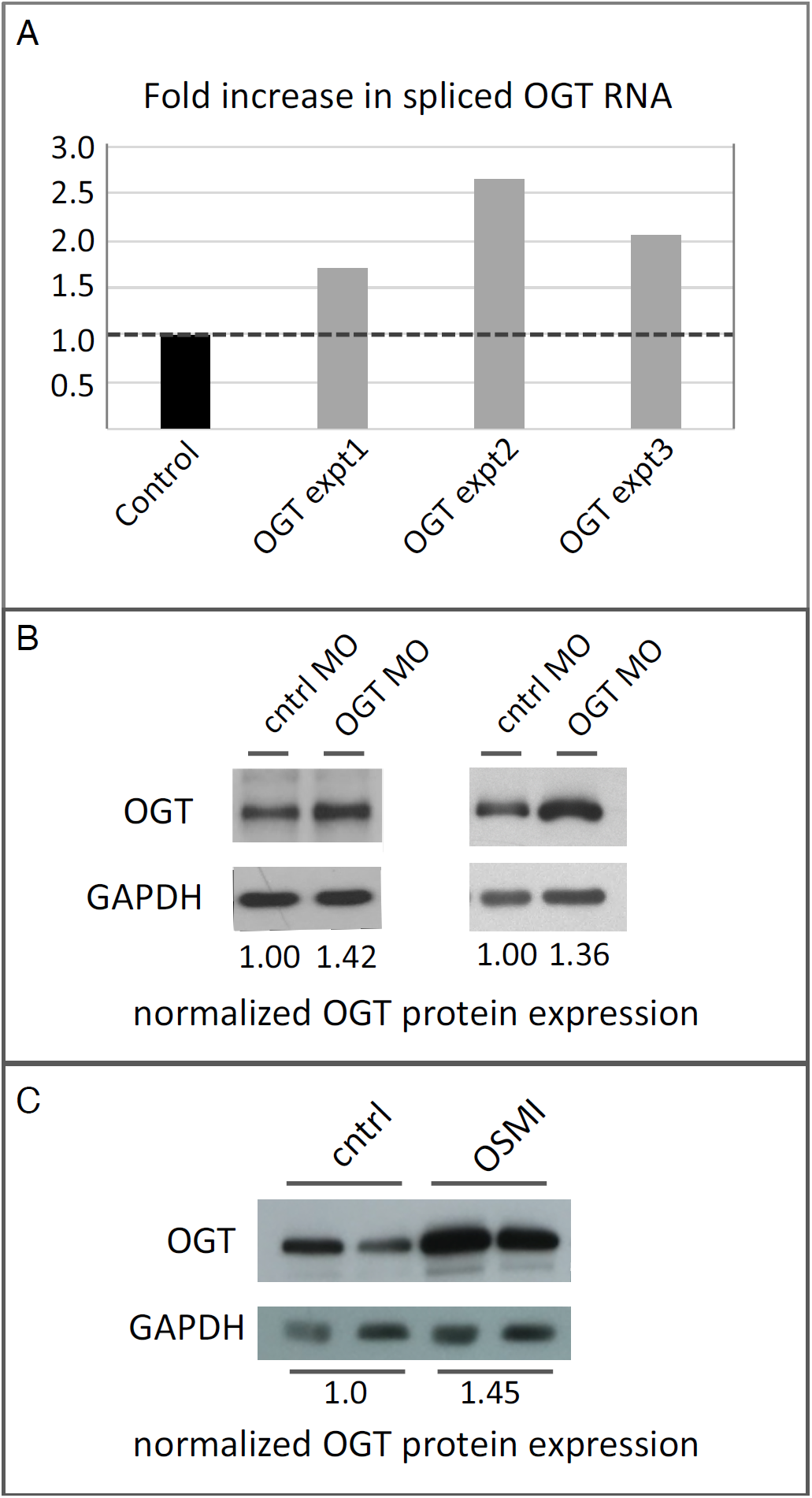
Decoy inhibition increases spliced RNA and protein output. A. Expression of spliced OGT transcript in decoy-inhibited cells, relative to expression in cells treated with a negative control MO. B. OGT protein expression in two independent experiments was increased in cells treated with the OGT 5’ decoy MO, compared to cells treated with a control MO. GAPDH expression was used to normalize protein loading. In both experiments inhibition of IR was accompanied by ∼1.4-fold increase in protein expression. C. OGT protein expression in two independent experiments was increased in cells treated with the OGT inhibitor OSMI-1, compared to cells treated with buffer alone.

Finally, we assessed erythroblast OGT protein expression as a function of variation in IR efficiency. Equal amounts of protein from control MO- or OGT MO-treated cells were immunoblotted with anti-OGT antibodies (Figure 5B). Densitometric analysis of OGT expression, normalized to expression of a control protein, GAPDH, revealed ∼1.4-fold increase in OGT protein levels in two independent experiments (compare control lanes to OGT MO lanes). Interestingly, the effects of decoy-targeting MOs on OGT protein expression in erythroblasts were similar to the effects induced by pharmacological treatment with the OGT inhibitor OSMI1 (Figure 5C). Similar effects of OSMI1 treatment on OGT expression were reported earlier in cancer cell lines (Park et al. 2017).

## Discussion

The decoy exon model proposes that intron retention levels can be controlled by modulating the balance between two competing splice site interactions: (1) productive cross-intron interactions, involving annotated splice sites at the intron termini, that promote splicing catalysis to remove the intron, and (2) nonproductive interactions, involving contacts between internal decoy exon(s) and intron terminal splice sites, that function mainly to block intron excision and promote intron retention. The latter are spliced inefficiently or not at all, presumably due to arrest of spliceosomal assembly at a pre-catalytic complex by mechanisms yet to be explored. This study validates a major prediction of the decoy hypothesis, namely, that blocking decoy exon function in endogenous pre-mRNA should shift RNA processing in favor of better intron removal. All four genes targeted with decoy-specific antisense MOs were shown to exhibit substantial decreases in IR. Interestingly, for two genes (SF3B1 and SPTA1), blocking the decoy exon essentially eliminated IR, suggesting that the decoy pathway may be the sole determinant of IR. In contrast, IR was not completely abrogated by antisense treatment in the OGT gene, consistent with the co-existence of decoy-independent IR mechanisms (Monteuuis et al. 2019; Braun et al. 2017; Cho et al. 2014; Wong et al. 2017). Finally, in the one case tested, OGT, decreased IR was accompanied by increases in spliced RNA and protein expression.

The IR transcripts studied here regulate expression of genes with diverse roles in erythropoiesis. Three function in general biochemical processes such as O-GlcNAc homeostasis (OGT) and pre-mRNA splicing (SF3B1 and SNRNP70), that are widely important in both erythroid and nonerythroid cells. Presumably the decoy mechanism actively regulates these genes in many different cell types. In contrast, SPTA1 functions predominantly in erythroid cells where it encodes a major structural component of the membrane skeleton that mechanical supports the eventual red cell membrane. Given the measured PIR values of 25-75% in these genes, full inhibition of IR could lead to 1.3-4-fold increases in protein expression, with most genes capable of ≤2-fold changes based on these bulk measures of IR. We speculate that the major purpose of decoy-mediated IR may be fine tuning of expression according to the cell’s physiological needs. In fact, IR has already been shown to tune OGT expression in cancer cell lines via an intronic element (Park et al. 2017) that likely operates via the decoy mechanism. For SPTA1, decoy-mediated intron retention could function in a similar manner to balance expression of the alpha and beta spectrin chains, two high molecular weight proteins that form an extended heterodimer that assembles into higher order structures supporting the red cell membrane. An imbalance of spectrin chains might be detrimental to human erythroblasts, and control of IR could serve to equalize the cellular content of these binding partner proteins. Finally, differentiating erythroblasts might dynamically regulate IR for splicing factor genes so that RNA splicing capacity could adapt to changes pre-mRNA abundance as thousands of genes are down-regulated during terminal erythropoiesis (An et al. 2014).

Interestingly, global comparison of RNA and protein abundance profiles in differentiating human erythroblasts has revealed discordant expression patterns that can be explained in part by IR (Gautier et al. 2016). As cells progressively differentiate into late stage erythroblasts, profiling experiments have shown that genes displaying increased RNA levels but decreased protein expression are enriched in IR-transcripts. In such cases IR may function to down-regulate productive gene expression in a post-transcriptional manner. We propose that decoy-mediated IR contributes substantially to this phenomenon, since erythroblasts express an estimated 400 retained introns embedded with candidate decoy exon(s) (Parra M. et al. 2018). Moreover, the number of functional decoys could be greater, because many of intronic U2AF binding sites detected in K562 cells do not align with splice junction-predicted erythroblast decoy exons. These U2AF sites of unknown function, perhaps regulated by novel RBP cofactors (Sutandy et al. 2018), could represent ‘silent’ decoys that promote IR without ever being catalytically spliced. Preliminary experiments supporting this idea are under further investigation.

The discovery of decoy exons provides new evidence that many unannotated splicing elements reside in deep intron space, hundreds to many thousands of nucleotides from the regulated splice sites, and that they play essential roles in regulating proper splice patterns via several mechanisms (Ule and Blencowe 2019). The decoy model discussed here is presumably employed in diverse cell types, since many decoy-containing introns are retained in a wide range of non-erythroid cell types. Another mechanism dependent on deep intron splicing elements is recursive splicing (RS). RS involves functional recognition of RS-exons, embedded deep within long introns, as critical splicing intermediates (Sibley et al. 2015; Joseph et al. 2018). Finally, a mechanism termed intrasplicing requires deep intron splicing elements, located tens of kilobases upstream of the regulated splice acceptors, to promote nested splicing reactions required for proper splice site selection in two paralogs of the protein 4.1 gene family (Parra M. K. et al. 2008; Parra M. K. et al. 2012). In various contexts, antisense oligonucleotides that block deep intron elements have been used to demonstrate their functional importance in splicing of endogenous pre-mRNAs (Parra M. K. et al. 2012; Sibley et al. 2015).

Finally, the current results suggest future clinical applications of antisense reagents for the purpose of improving gene expression. There is substantial evidence that deep intron mutations can cause human disease (Vaz-Drago et al. 2017), and increasing precedence for the use of antisense oligonucleotides to block intronic elements that restrict productive splicing so as to improve gene expression. As one example, antisense reagents that block intronic splicing silencer(s) downstream of SMN2 exon 7 improve productive splicing, and in fact an antisense oligonucleotide drug has already been approved for therapy of patients with spinal muscular atrophy due to inactivation of SMN1 (Bennett et al. 2019). Antisense reagents can also restore correct splicing, and improve gene expression, by blocking deep intron splicing mutations that activate inclusion of cryptic noncoding exons, e.g., in the breast cancer gene BRCA2 (Anczukow et al. 2012) and the deaf-blindness gene USH2A (Slijkerman et al. 2016). Our data show that antisense oligonucleotides can in principle increase protein expression by blocking decoy splice sites in retained introns, which could allow an intact allele to increase protein output to compensate for deficiencies caused by mutational inactivation of disease alleles.

## Materials and Methods

### Erythroblast culture

CD34+ erythroid progenitors were enriched from cord blood and cultured under conditions previously shown to support selective growth and differentiation of erythroid cells (Hu et al. 2013). For electroporation, 106 erythroblasts at day 11 of culture were electroporated at room temperature in supplemented P3 solution using a Lonza 4D-Nucleofector system with the ER 100 pulse code. 25nt morpholinos, antisense to the regions highlighted in Figure 1B, were obtained from Gene Tools LLC (Philomath, OR), maintained in sterile saline solution, and added to the cells at 30*μ*M final concentration just prior to electroporation. After electroporation cells were incubated in culture medium at 37°C for 2 days before further processing. When RNA and protein were isolated from the same sample, ∼2.5×105 cells were used for the RNA preparation and 7.5×105 cells for protein purification. Morpholino sequences antisense to the 5’ splice sites of the targeted decoys were as follows:

OGT, 5’-gtggcagttacaaac|ccgttac|CAT-3’;

SPTA1, 5’-ctggctggaac|ctcttac|GTGGCTG-3’;

SF3B1, 5’-atccggaatacgtac|ACTTTCGTGC-3’;

SNRNP70, 5’-ccatgatacaaac|ccttataccaac|-3’.

Sequences antisense to the intron are in lower case; sequences antisense to the exon are in upper case; EXON|intron boundaries are marked by vertical lines.

### RT-PCR

Total RNA was extracted from human erythroid progenitors and analyzed by standard RT-PCR methods as described (Pimentel et al. 2016). For quantitative analysis, RT-qPCR was performed using an Applied Biosystems 7500 Fast Real-Time PCR System with Quanta SYBR Green Fastmix low ROX reagents. The SYBR Green buffers were supplemented with forward and reverse primers at a final concentration of 0.5*μ*M, and the spliced RNA or IR-RNA DNAs amplified using the following program: initial denaturation (1 cycle): 94°C for 15 min; amplification stage (40 cycles): 95°C for 10 sec, 60°C for 25 sec, and 72°C for 30 sec; final extension at 72°C for 30 sec. Size and identity of qPCR products were confirmed by gel electrophoresis (Figure S1) and by DNA sequencing. The relative expression of each gene was calculated using the comparative ΔCt method after normalizing to the ACTB control.

### Western blot analysis

After electroporation followed by an additional ∼48hrs of culture, an estimated 7.5×105 cells were pelleted and stored at -80°C. Protein was subsequently isolated from lysed cells, subjected to SDS-polyacrylamide gel electrophoresis, and immunoblotted using rabbit polyclonal antibody against OGT (Proteintech group, Inc., Rosemont, IL; cat. no. 11576-2) at 1:4000 dilution, or rabbit polyclonal antibody against GAPDH (Sigma, cat.no. G9545) at 1:10,000 dilution.

## Acknowledgments

This work was funded by National Institutes of Health grant 5R01DK108020 (J.G.C.) and by the Director, Office of Science and Office of Biological & Environmental Research of the US Department of Energy (DE-AC02-05CH1123).

## Figures

**Table 1.**
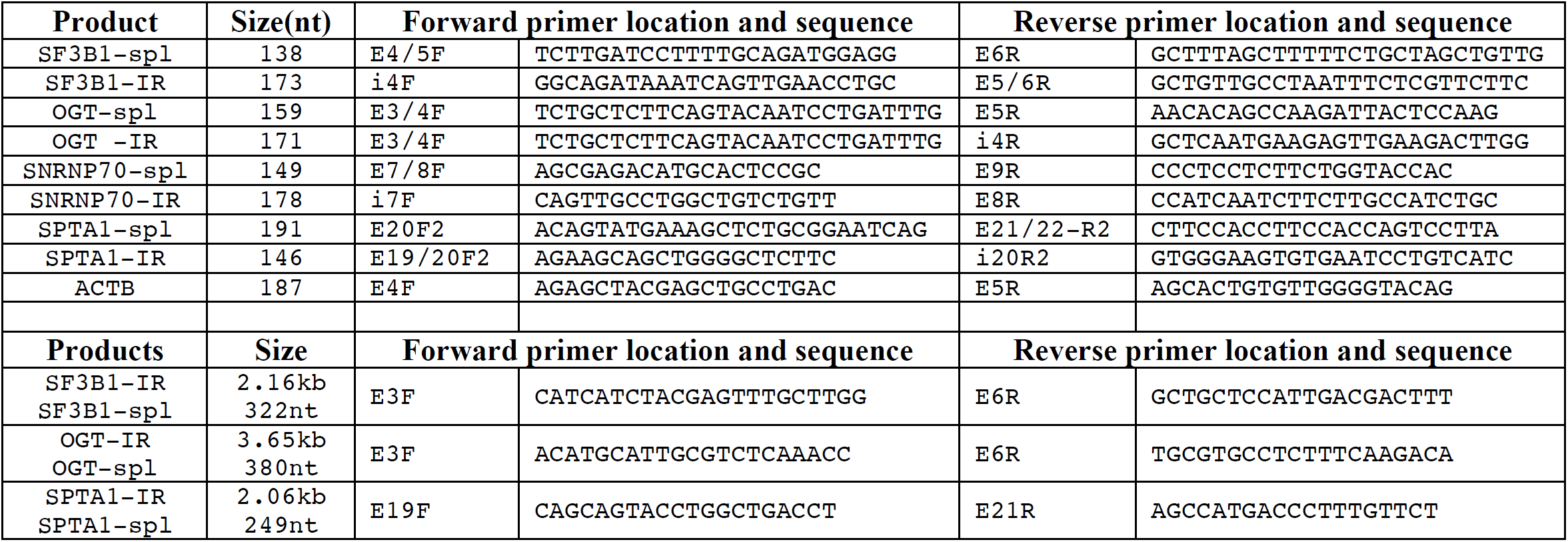
Primers used for PCR.

**Figure S1.**
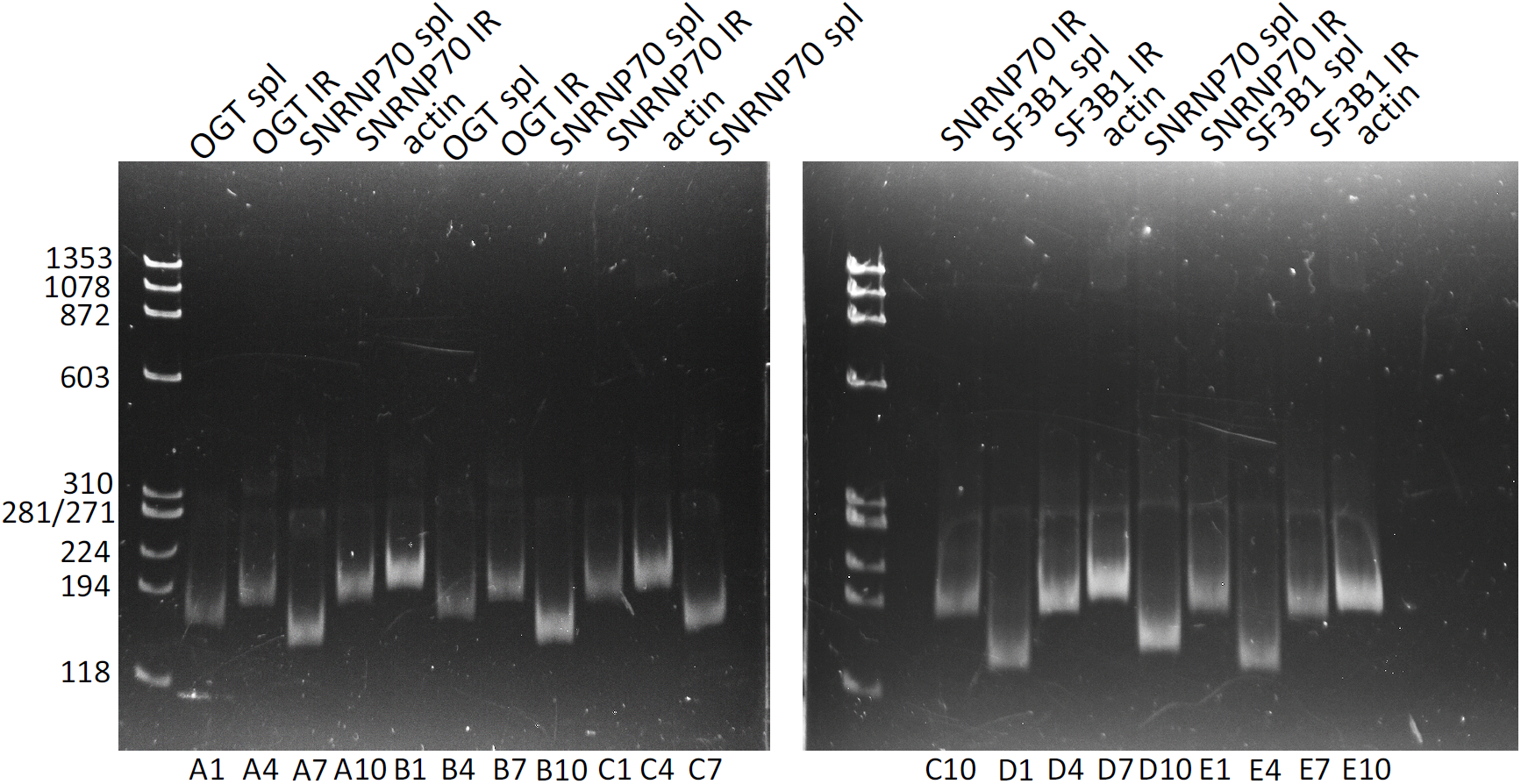
Gel analysis of qPCR products. Amplification products representing each of the major qPCR reactions for spliced (spl) and IR-transcripts (IR) were analyzed by polyacrylamide gel electrophoresis. Designations below the gel indicate well numbers from the 96 well plate used for amplification. Numbers at the left indicate DNA size standards in nucleotides.

## References

Adusumalli, S., Z. K. Ngian, W. Q. Lin, T. Benoukraf and C. T. Ong (2019). Increased intron retention is a post-transcriptional signature associated with progressive aging and Alzheimer’s disease. Aging Cell 18: e12928.

An, X., V. P. Schulz, J. Li, K. Wu, J. Liu, F. Xue, J. Hu, N. Mohandas and P. G. Gallagher (2014). Global transcriptome analyses of human and murine terminal erythroid differentiation. Blood 123: 3466–3477.

Anczukow, O., M. Buisson, M. Leone, C. Coutanson, C. Lasset, A. Calender, O. M. Sinilnikova and S. Mazoyer (2012). BRCA2 deep intronic mutation causing activation of a cryptic exon: opening toward a new preventive therapeutic strategy. Clin Cancer Res 18: 4903–4909.

Bennett, C. F., A. R. Krainer and D. W. Cleveland (2019). Antisense Oligonucleotide Therapies for Neurodegenerative Diseases. Annu Rev Neurosci 42: 385–406.

Boothby, T. C., R. S. Zipper, C. M. van der Weele and S. M. Wolniak (2013). Removal of retained introns regulates translation in the rapidly developing gametophyte of Marsilea vestita. Dev Cell 24: 517–529.

Boutz, P. L., A. Bhutkar and P. A. Sharp (2015). Detained introns are a novel, widespread class of post-transcriptionally spliced introns. Genes Dev 29: 63–80.

Braun, C. J., M. Stanciu, P. L. Boutz, J. C. Patterson, D. Calligaris, F. Higuchi, R. Neupane, S. Fenoglio, D. P. Cahill, H. Wakimoto, et al. (2017). Coordinated Splicing of Regulatory Detained Introns within Oncogenic Transcripts Creates an Exploitable Vulnerability in Malignant Glioma. Cancer Cell 32: 411–426 e411.

Braunschweig, U., N. L. Barbosa-Morais, Q. Pan, E. N. Nachman, B. Alipanahi, T. Gonatopoulos-Pournatzis, B. Frey, M. Irimia and B. J. Blencowe (2014). Widespread intron retention in mammals functionally tunes transcriptomes. Genome Res 24: 1774–1786.

Brugiolo, M., V. Botti, N. Liu, M. Muller-McNicoll and K. M. Neugebauer (2017). Fractionation iCLIP detects persistent SR protein binding to conserved, retained introns in chromatin, nucleoplasm and cytoplasm. Nucleic Acids Res 45: 10452–10465.

Chen, L., R. Weinmeister, J. Kralovicova, L. P. Eperon, I. Vorechovsky, A. J. Hudson and I. C. Eperon (2017). Stoichiometries of U2AF35, U2AF65 and U2 snRNP reveal new early spliceosome assembly pathways. Nucleic Acids Res 45: 2051–2067.

Cho, V., Y. Mei, A. Sanny, S. Chan, A. Enders, E. M. Bertram, A. Tan, C. C. Goodnow and T. D. Andrews (2014). The RNA-binding protein hnRNPLL induces a T cell alternative splicing program delineated by differential intron retention in polyadenylated RNA. Genome Biol 15: R26.

Dvinge, H. and R. K. Bradley (2015). Widespread intron retention diversifies most cancer transcriptomes. Genome Med. 7: 45.

Edwards, C. R., W. Ritchie, J. J. Wong, U. Schmitz, R. Middleton, X. An, N. Mohandas, J. E. Rasko and G. A. Blobel (2016). A dynamic intron retention program in the mammalian megakaryocyte and erythrocyte lineages. Blood 127: e24–e34.

Gautier, E. F., S. Ducamp, M. Leduc, V. Salnot, F. Guillonneau, M. Dussiot, J. Hale, M. C. Giarratana, A. Raimbault, L. Douay, et al. (2016). Comprehensive Proteomic Analysis of Human Erythropoiesis. Cell Rep 16: 1470–1484.

Hu, J., J. Liu, F. Xue, G. Halverson, M. Reid, A. Guo, L. Chen, A. Raza, N. Galili, J. Jaffray, et al. (2013). Isolation and functional characterization of human erythroblasts at distinct stages: implications for understanding of normal and disordered erythropoiesis in vivo. Blood 121: 3246–3253.

Jacob, A. G. and C. W. J. Smith (2017). Intron retention as a component of regulated gene expression programs. Hum Genet 136: 1043–1057.

Joseph, B., S. Kondo and E. C. Lai (2018). Short cryptic exons mediate recursive splicing in Drosophila. Nat Struct Mol Biol 25: 365–371.

Lovci, M. T., D. Ghanem, H. Marr, J. Arnold, S. Gee, M. Parra, T. Y. Liang, T. J. Stark, L. T. Gehman, S. Hoon, et al. (2013). Rbfox proteins regulate alternative mRNA splicing through evolutionarily conserved RNA bridges. Nat Struct Mol Biol 20: 1434–1442.

Luisier, R., G. E. Tyzack, C. E. Hall, J. S. Mitchell, H. Devine, D. M. Taha, B. Malik, I. Meyer, L. Greensmith, J. Newcombe, et al. (2018). Intron retention and nuclear loss of SFPQ are molecular hallmarks of ALS. Nat Commun 9: 2010.

Mauger, O., F. Lemoine and P. Scheiffele (2016). Targeted Intron Retention and Excision for Rapid Gene Regulation in Response to Neuronal Activity. Neuron 92: 1266–1278.

Monteuuis, G., J. J. L. Wong, C. G. Bailey, U. Schmitz and J. E. J. Rasko (2019). The changing paradigm of intron retention: regulation, ramifications and recipes. Nucleic Acids Res 47: 11497–11513.

Naro, C., A. Jolly, S. Di Persio, P. Bielli, N. Setterblad, A. J. Alberdi, E. Vicini, R. Geremia, P. De la Grange and C. Sette (2017). An Orchestrated Intron Retention Program in Meiosis Controls Timely Usage of Transcripts during Germ Cell Differentiation. Dev Cell 41: 82–93 e84.

Ni, T., W. Yang, M. Han, Y. Zhang, T. Shen, H. Nie, Z. Zhou, Y. Dai, Y. Yang, P. Liu, et al. (2016). Global intron retention mediated gene regulation during CD4+ T cell activation. Nucleic Acids Res 44: 6817–6829.

Ninomiya, K., N. Kataoka and M. Hagiwara (2011). Stress-responsive maturation of Clk1/4 pre-mRNAs promotes phosphorylation of SR splicing factor. J Cell Biol 195: 27–40.

Park, S. K., X. Zhou, K. E. Pendleton, O. V. Hunter, J. J. Kohler, K. A. O’Donnell and N. K. Conrad (2017). A Conserved Splicing Silencer Dynamically Regulates O-GlcNAc Transferase Intron Retention and O-GlcNAc Homeostasis. Cell Rep 20: 1088–1099.

Parra, M., B. W. Booth, R. Weiszmann, B. Yee, G. W. Yeo, J. B. Brown, S. E. Celniker and J. G. Conboy (2018). An important class of intron retention events in human erythroblasts is regulated by cryptic exons proposed to function as splicing decoys. RNA 24: 1255–1265.

Parra, M. K., T. L. Gallagher, S. L. Amacher, N. Mohandas and J. G. Conboy (2012). Deep intron elements mediate nested splicing events at consecutive AG-dinucleotides to regulate alternative 3’ splice site choice in vertebrate 4.1 genes. Mol Cell Biol 32: 2044–2053.

Parra, M. K., J. S. Tan, N. Mohandas and J. G. Conboy (2008). Intrasplicing coordinates alternative first exons with alternative splicing in the protein 4.1R gene. EMBO J 27: 122–131.

Pendleton, K. E., B. Chen, K. Liu, O. V. Hunter, Y. Xie, B. P. Tu and N. K. Conrad (2017). The U6 snRNA m(6)A Methyltransferase METTL16 Regulates SAM Synthetase Intron Retention. Cell 169: 824–835 e814.

Pendleton, K. E., S. K. Park, O. V. Hunter, S. M. Bresson and N. K. Conrad (2018). Balance between MAT2A intron detention and splicing is determined cotranscriptionally. RNA 24: 778–786.

Pimentel, H., M. Parra, S. Gee, D. Ghanem, X. An, J. Li, N. Mohandas, L. Pachter and J. G. Conboy (2014). A dynamic alternative splicing program regulates gene expression during terminal erythropoiesis. Nucleic Acids Res 42: 4031–4042.

Pimentel, H., M. Parra, S. L. Gee, N. Mohandas, L. Pachter and J. G. Conboy (2016). A dynamic intron retention program enriched in RNA processing genes regulates gene expression during terminal erythropoiesis. Nucleic Acids Res 44: 838–851.

Pirnie, S. P., A. Osman, Y. Zhu and G. G. Carmichael (2017). An Ultraconserved Element (UCE) controls homeostatic splicing of ARGLU1 mRNA. Nucleic Acids Res 45: 3473–3486.

Rekosh, D. and M. L. Hammarskjold (2018). Intron retention in viruses and cellular genes: Detention, border controls and passports. Wiley Interdiscip Rev RNA 9: e1470.

Shalgi, R., J. A. Hurt, S. Lindquist and C. B. Burge (2014). Widespread inhibition of posttranscriptional splicing shapes the cellular transcriptome following heat shock. Cell Rep 7: 1362–1370.

Sibley, C. R., W. Emmett, L. Blazquez, A. Faro, N. Haberman, M. Briese, D. Trabzuni, M. Ryten, M. E. Weale, J. Hardy, et al. (2015). Recursive splicing in long vertebrate genes. Nature 521: 371–375.

Slijkerman, R. W., C. Vache, M. Dona, G. Garcia-Garcia, M. Claustres, L. Hetterschijt, T. A. Peters, B. P. Hartel, R. J. Pennings, J. M. Millan, et al. (2016). Antisense Oligonucleotide-based Splice Correction for USH2A-associated Retinal Degeneration Caused by a Frequent Deep-intronic Mutation. Mol Ther Nucleic Acids 5: e381.

Sutandy, F. X. R., S. Ebersberger, L. Huang, A. Busch, M. Bach, H.-S. Kang, J. Fallmann, D. Maticzka, R. Backofen, P. F. Stadler, et al. (2018). In vitro iCLIP-based modeling uncovers how the splicing factor U2AF2 relies on regulation by cofactors. Genome Research 28: 699–713.

Ule, J. and B. J. Blencowe (2019). Alternative Splicing Regulatory Networks: Functions, Mechanisms, and Evolution. Mol Cell 76: 329–345.

Vaz-Drago, R., N. Custodio and M. Carmo-Fonseca (2017). Deep intronic mutations and human disease. Hum Genet 136: 1093–1111.

Wong, J. J., D. Gao, T. V. Nguyen, C. T. Kwok, M. van Geldermalsen, R. Middleton, N. Pinello, A. Thoeng, R. Nagarajah, J. Holst, et al. (2017). Intron retention is regulated by altered MeCP2-mediated splicing factor recruitment. Nat Commun 8: 15134.

Wong, J. J., W. Ritchie, O. A. Ebner, M. Selbach, J. W. Wong, Y. Huang, D. Gao, N. Pinello, M. Gonzalez, K. Baidya, et al. (2013). Orchestrated intron retention regulates normal granulocyte differentiation. Cell 154: 583–595.

